# PAKC: A novel Panel of HLA class I Antigen presentation machinery Knockout Cells from the same genetic origin

**DOI:** 10.1101/2020.01.24.917807

**Authors:** Antonius A. de Waard, Tamara Verkerk, Marlieke L.M. Jongsma, Kelly Hoefakker, Sunesh Sethumadhavan, Carolin Gerke, Sophie Bliss, George M.C. Janssen, Arnoud H. de Ru, Frans H.J. Claas, Arend Mulder, Robert Tampé, Peter A. van Veelen, Anne Halenius, Robbert M. Spaapen

## Abstract

With the emergence of immunotherapies, the understanding of functional HLA class I antigen presentation to T cells is more relevant than ever. Current knowledge on antigen presentation is based on decades of research in a wide variety of cell types with varying antigen presentation machinery (APM) expression patterns, proteomes and HLA haplotypes. This diversity complicates the establishment of individual APM contributions to antigen generation, selection and presentation. Therefore, we generated a novel Panel of APM Knockout Cell lines (PAKC) from the same genetic origin. After CRISPR/Cas9 genome-editing of ten individual APM components in a human cell line, we derived clonal cell lines and confirmed their knockout status and phenotype. We then show how PAKC will accelerate research on the functional interplay between APM components and their role in antigen generation and presentation. This will lead to improved understanding of peptide-specific T cell responses in infection, cancer and autoimmunity.

**Key points:**

1. We generated a panel of cell lines to study HLA class I antigen presentation
2. We show how this will spark research in infection, tumor biology and autoimmunity

## Introduction

HLA class I (HLA-I) proteins are crucial for the onset of T cell responses during infections, cancer or autoimmunity(1–3). HLA-I continuously samples peptides from intracellularly degraded proteins and presents these on the cell surface to CD8^+^ T cells(3). To escape T cell surveillance, various viruses express inhibitors of the machinery required for antigen processing and loading onto HLA-I (antigen presentation machinery; APM)(2). For the same reason, tumor cells acquire genetic impairments in the APM, especially after immunotherapy(4). On the other hand, HLA-I alleles and functional polymorphisms in the APM have been highly associated with the development of several auto-inflammatory diseases(1). In the past decades, the individual function of each of the APM components has been widely studied(3). These studies revealed that newly synthesized HLA-I heavy chains are *N*-glycosylated and consecutively bound by the lectin chaperones calnexin (CNX) and calreticulin (CALR). Several glycosidases, such as glucosidase IIα, trim the glycan tree generating the core substrate for complex *N*-glycosylation later on in the Golgi. After recruiting beta-2 microglobulin (B2M), the HLA-I heavy chain associates with the peptide loading complex consisting of ERp57, tapasin and the transporter associated with peptide loading (TAP1/TAP2 heterodimer)(5). The TAP heterodimer transports peptides derived from degraded proteins from the cytosol into the ER lumen, where they will be occasionally trimmed by the aminopeptidase ERAP1. Tapasin and ERp57 assist in the binding of peptides into the so-called HLA-I peptide binding groove, after which the mature HLA-I heterotrimer will be shuttled via the Golgi to the plasma membrane to present its peptide to CD8^+^ T cells(6).

These studies were conducted using various (sets of) murine or human model cell lines with varying APM expression patterns, proteomes and HLA haplotypes, complicating the comparison of their individual contributions to antigen processing and presentation. Moreover, the molecular interplay between the different APM components may have confounding effects in assays with single APM component mutant cell lines. Previous efforts have established 721 lymphoblastic cell lines with gamma irradiation-induced TAP1, tapasin and HLA-I mutants, which have been used extensively to enlarge the knowledge on these factors. To address the current questions in the field, it is essential to generate a broader and more defined model system capable of overcoming these issues and addressing the current questions in the field. Here, we generated a novel panel of HLA-I APM knockout cell lines, designated PAKC, on the background of human HAP1 cells. Using CRISPR/Cas9, we generated ten clonal cell lines knocked out for individual components of the HLA-I antigen processing and presentation pathway. We illustrate how the availability of this collection to the community will spark state-of-the-art research to explain associations of the APM and certain HLA-I alleles with diseases. Additionally, we show the opportunities of PAKC to improve the rules of peptide processing and presentation which will further increase our understanding of infection, cancer and autoimmunity.

## Materials and Methods

### Cell culture

HAP1 and HEK293T cell lines were cultured at 37 °C and 5% CO_2_ in IMDM (Gibco) supplemented with 10% FCS and antibiotics (PenStrep; Invitrogen). CD8^+^ T cell clones recognizing peptides derived from the endogenously expressed proteins USP11 and SSR1(7, 8) were expanded using a standard feeder mix in IMDM supplemented with 5% human serum (Sanquin) and 5% FCS(9).

### Genome editing and overexpression

gRNAs used for gene editing are listed in Supplementary Table I. pX458 containing a B2M targeting gRNA was kindly provided by Dr. R. Mezzadra. ERp57 and tapasin targeting gRNAs in pX330 were co-transfected with a Blasticidin S-resistance overexpression vector as previously described(10). A gRNA targeting a conserved region in HLA-A, -B, -C and -G in a pX330 vector was transfected into HAP1 cells using X-tremeGENE (Roche). Single cells were FACS sorted based on W6/32 negativity to obtain knockout clones for HLA-A, -B and -C. gRNAs targeting CNX, ERAP1, CALR and glucosidase IIα in pLentiCRISPRv2 and a gRNA targeting Tap1 in pL.CRISPR.efs.GFP were co-transfected with packaging plasmids psPAX2 and pVSVg, and pAdVAntage (Promega) using polyethylenimine (PEI; Polyscience) into HEK293T cells for virus production. puc2CL6IP HLA-B*27:05/09 constructs (kind gift of Dr. S. Springer(11)) were co-transfected using PEI with packaging plasmids pVSVg and NLBH into HEK293T cells for virus production. Viral supernatants of other HLA alleles contained a ΔNGFR marker gene (kind gift of Dr. M Griffioen) (12). Viral supernatant was filtered and used for transduction by spinoculation in the presence of 8 μg/mL protamine sulfate. Several genome-edited cells were selected using puromycin or blasticidin S, before limiting dilution cloning. TAP1 knockout cells were enriched by FACS sort on GFP positivity.

### Sanger sequencing

Genomic DNA was isolated using the NucleoSpin Tissue kit (Machery-Nagel). DNA was amplified and sequenced using BigDye v1.1 (Applied Biosystems) and specific primers listed in Supplementary Table II.

### Immunoblotting

Proteins were separated by SDS-PAGE (Bis-Tris NuPAGE 12% gel (CALR immunoblot, 10% Bis-Tris NuPAGE gel), ThermoFisher) and transferred to a PVDF membrane (Amersham Hybond-P, GE Healthcare) (CALR, nitrocellulose, Invitrogen) at 25 V for 12 min by using the semidry Trans-Blot Turbo (BioRad) system (CALR, iBlot2 semi-dry blotting system, Invitrogen). Membranes were blocked for 1 h with 3% (w/v) non-fat milk powder in PBS/0.1% (v/v) Tween (PBST) (CALR, 10% Roche WBR in PBST) and incubated with primary antibody (Supplementary Table III) over night (CALR, 1h) at 4 °C. After washing three times in PBST, the membrane was incubated with secondary antibody for 1 h RT and washed again in PBST. Blots were incubated with Clarity Western ECL reagent (BioRad) (CALR, Pierce ECL plus, ThermoFisher Scientific), the signal was detected with a Lumi-Imager (Vilber, Fusion FX) (CALR, Image Lab software, Bio-Rad) and analyzed with Evolution-Capt Edge software.

### FACS

Cells were incubated with specific antibodies (Supplementary Table III) diluted in PBS for 30 min on ice. Unconjugated antibodies were washed away in three times before secondary stain. Stained cells were sorted on a BD Aria II or fixed in PBS/1% (v/v) formaldehyde/1 μM DAPI (Sigma-Aldrich) and analyzed by BD flow cytometers (Fortessa or LSR II). Data was analyzed using FlowJo (Tree Star, Inc). Data were gated on DAPI negativity, single cells, time and if applicable an ΔNGFR reporter.

### IFN-γ stimulation

Cells were cultured with 0, 10, 20, 40 or 80 U/mL IFN-γ (PeproTech) for 1 or 2 d before analysis by qPCR and FACS, respectively.

### qPCR

RNA extraction and qPCR was performed as described previously(13). Gene expression was determined using SYBR green and the StepOnePlus (ThermoFisher). Quantification was done using the ΔΔC_T_ method. 18S rRNA expression was used as an internal reference, primers are listed in Supplementary Table II.

### Pulse-chase

Cells were incubated for 24 h with 100 U/mL IFN-γ (BioLegend), pulse-chase was performed as previously described(14). Cells were metabolically labeled with 0.2 mCi/mL for 30 min, lysed and cleared from membrane debris at 16,200 xg for 30 min at 4 °C. Lysates were incubated with W6/32 or MaP.ERp57 (Abcam) for 1 h at 4 °C in an overhead tumbler before retrieving immune complexes with protein G Sepharose (GE Healthcare). Beads were washed and complexes were treated with Endoglycosidase H (New England Biolabs) according to manufacturer’s protocol. Prior to loading on a 10-14% gradient SDS-PAGE gel, complexes were dissociated at 95 °C in sample buffer (150mM DTT).

### Mass spectrometry of HLA-class I peptides

335 ×10^6^ cells were lysed in 3.3 ml lysis buffer and processed as described(15). Peptides were lyophilized, dissolved in 95/3/0.1% (v/v/v) water/acetonitrile/formic acid (WAFA) and analyzed by on-line C18 nanoHPLC MS/MS with a system consisting of an Ultimate3000nano gradient HPLC system (Thermo, Bremen, Germany) and an Exploris480 mass spectrometer (Thermo). Fractions were injected onto a cartridge precolumn (300 μm × 5 mm, C18 PepMap, 5 μm, 100 A) and eluted via a homemade analytical nano-HPLC column (50 cm × 75 μm; Reprosil-Pur C18-AQ 1.9 μm, 120 A (Dr. Maisch, Ammerbuch, Germany)). The gradient was run from 2% to 36% (v/v) solvent B (20/80/0.1 (v/v/v) WAFA) in 120 min. The nano-HPLC column was drawn to a tip of ~10 μm and acted as the electrospray needle of the MS source. The MS was operated in data-dependent MS/MS mode for a cycle time of 3 s, with an HCD collision energy at 30 V and recording of the MS2 spectrum in the orbitrap, with a quadrupole isolation width of 1.6 Da. In the master scan (MS1) the resolution was 60,000, the scan range 300-1,500, at an AGC target of 1,000,000 at maximum fill time of 120 ms. A lock mass correction on the background ion m/z=445.12 was used. Precursors were dynamically excluded after n=1 with an exclusion duration of 45 s, and with a precursor range of 20 ppm. Charge states 1-3 were included. For MS2, the first mass was set to 110 Da, and the MS2 scan resolution was 30,000 at an AGC target of 100,000 at maximum fill time of 120 ms.

Raw data were first converted to peak lists using Proteome Discoverer version 2.2 (Thermo Electron), and submitted to the Uniprot Homo sapiens minimal database (20,205 entries), using Mascot v. 2.2.04 (www.matrixscience.com) for protein identification (10 ppm precursor, 0.02 Da deviation of fragment mass, no enzyme specified). Methionine oxidation and cysteinylation on cysteine were set as a variable modification. Peptides with an FDR<1% in combination with a mascot ion score >35 were accepted.

Peptide affinities were predicted using NetMHC4.0 for HLA-A*02:01, HLA-B*40:01 and HLA-C*03:03, which has a comparable binding motif as the expressed HLA-C*03:04(16). Peptide sequence clustering was performed for all 9-mers using GibbsCluster-2.0 and Seq2Logo.

### T cell assays

Target cells were incubated with T cells in a 1:1 ratio for 18 h as previously described(17). IFN-γ or GM-CSF release in the cell culture supernatant was measured using ELISA according to the manufacturer’s protocol (Sanquin and BioLegend, respectively).

### Statistical analysis

Statistical testing was done by a one-way ANOVA followed by Dunnett’s multiple comparisons test (GraphPad Prism).

## Results and Discussion

We selected HAP1 as the parental cell line for creating PAKC as it has excellent cloning capacity, a high proliferation rate and a functional HLA-I antigen presentation pathway. Using CRISPR/Cas9 we created knockout cell lines for the currently known players in HLA-I antigen presentation: HLA-I heavy chain, CNX, glucosidase IIα, B2M, CALR, tapasin, ERp57, TAP1, TAP2 and ERAP1 (Figure 1A). ERAP2 and TABPBR were not targeted since RNA-seq data show that these are not expressed in HAP1 cells(18). Over time, we used different strategies to introduce the Cas9 and gRNAs targeting the coding sequence close to the start codon of each gene to create the individual knockout cell lines. First, tapasin and ERp57 knockout cell lines were created by gRNA-directed genomic insertion of a blasticidin S-resistance gene. The second approach consisted of gRNA-driven induction of random deletions or insertions to cause frameshifts in the genes encoding HLA-I, CNX, B2M, CALR, TAP1, TAP2, ERAP1 and glucosidase IIα. HLA-I was targeted using a single gRNA recognizing a conserved sequence in the HLA-A, -B and -C genes. Of note, this gRNA is also specific for HLA-G and has one or multiple mismatches with HLA-E and HLA-F genes.

**Figure 1.**
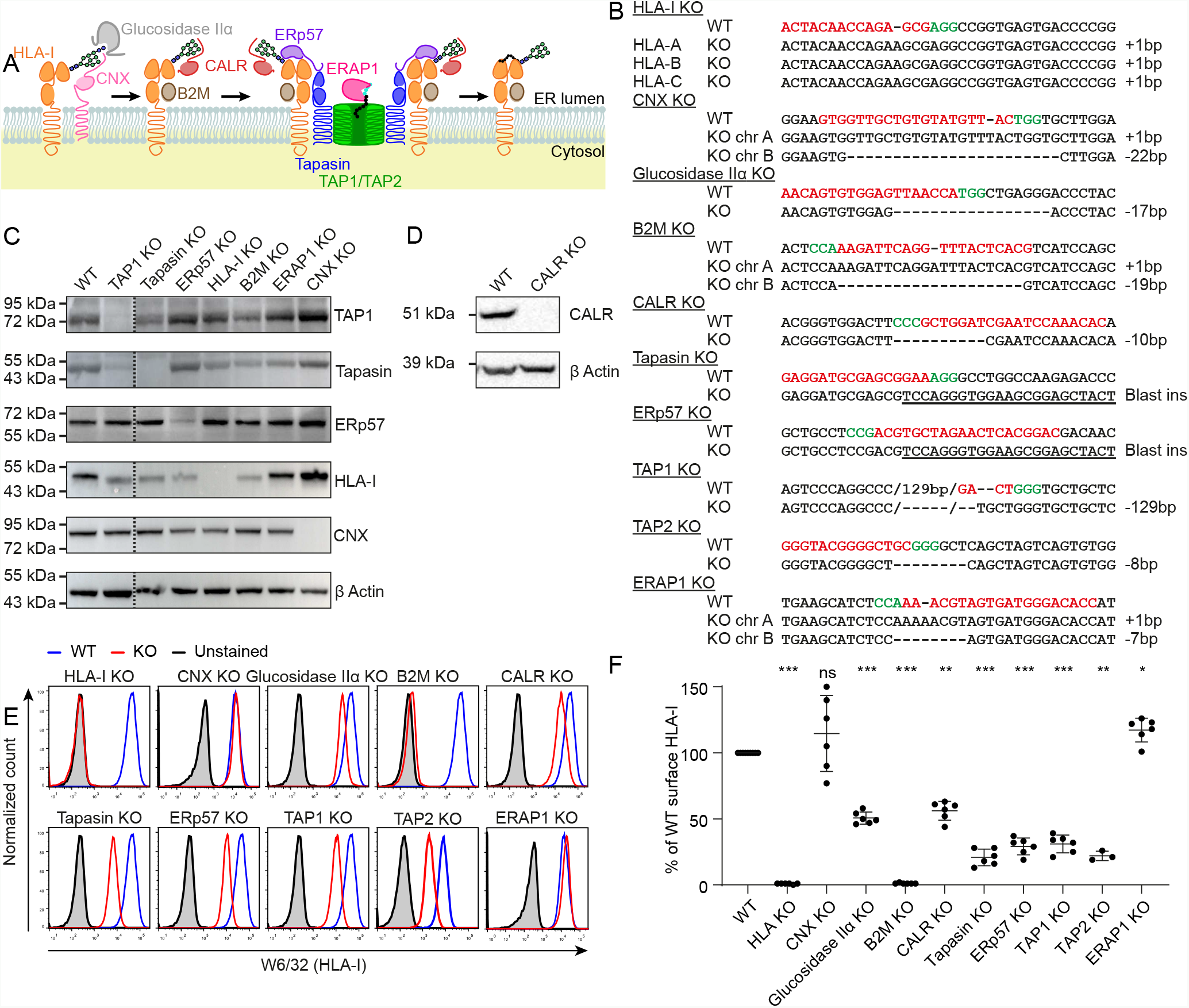
Validation of PAKC on sequence and protein level. (**A**) Schematic overview of the HLA-I antigen presentation pathway. (**B**) Sequence summary of one selected clone per targeted APM component. Shown are CRISPR/Cas9 induced mutations (dashes), the gRNA (red) and PAM sequence (green). Tapasin ERp57 knockout clones were generated by genomic insertion of a blasticidin S-resistance gene (underlined). Chr, chromosome. Raw data in Supplementary Figure 1. (**C**), (**D**) Immunoblots of knockout cell lines with indicated antibodies. (**E**), (**F**) FACS analysis of indicated knockout cells using the pan-HLA-I antibody W6/32. (**E**) Representative plots with wild type in blue, knockout in red and unstained control in grey. (**F**) Relative quantification, n=6 independent experiments (TAP2 knockout n=3). One-Way ANOVA before normalization, *p<0.05, **p<0.01, ***p<0.0001, ns=not significant. WT, wild type; KO, knockout.

After initial selection by antibiotics or FACS sort for a reporter, clonal cell lines were generated by limiting dilution and targeted gene regions were Sanger sequenced to evaluate loss of gene integrity. We then selected knockout lines for all target genes. Notably, the most detrimental genotype for TAP1 knockout clones was a deletion of 129 bp covering the entire second transmembrane domain (Figure 1B and Supplementary Figure 1)(19). Immunoblot analysis of this clone confirmed a complete lack of TAP1 expression. Similarly, we validated the absence of tapasin, ERp57, HLA-I heavy chain, CNX and CALR proteins in the respective knockout cell lines (Figure 1C and D).

As a final validation of PAKC, we characterized the effect of each knockout on HLA-I surface expression by flow cytometry using the HLA-I antibody W6/32 (Figure 1E and F). Except CNX and ERAP1, all APM knockouts induced a significant decrease of HLA-I surface expression confirming findings of others. ERAP1 knockout cells show a slight but significant increase in surface HLA-I, which has been observed before(20). The chaperones CALR and CNX are known to execute similar functions(21). Redundancy between these chaperones is a potential explanation for the absence of a phenotype in cells depleted of CNX. Inversely, CNX is incapable of fully substituting CALR function as HLA-I surface levels are decreased in CALR knockout cells. In depth studies into the mutual effect of these proteins on antigen presentation may thus require the additional generation of CNX and CALR double knockout cells.

We next evaluated whether PAKC provides also other opportunities to improve our understanding of HLA-I antigen presentation. In fact, our straightforward immunoblot analyses indicated that the knockout of TAP1, HLA-I, B2M or ERAP1 decreased tapasin expression, suggesting a novel role for each of these in stabilizing tapasin (Figure 1C). This is relevant to the ongoing debate whether the protease ERAP1 is accommodated within the peptide loading complex(22). The regulation of tapasin homeostasis and the functional consequences thereof are still unknown. In the opposite direction, tapasin has been described to stabilize TAP, showing the complexity of peptide loading complex regulation(23). These findings demonstrate that PAKC provides opportunities to understand the interplay between different APM components.

HLA-I is highly polymorphic with more than 17,000 different alleles identified in the human population. Several of these alleles and other APM polymorphisms have been associated with susceptibility or resistance to infection, the development of cancer and autoimmunity(1). To investigate the feasibility of studying such disease-associated variations with PAKC, we reconstituted a set of ten HLA-I alleles into wild type HAP1 cells. These alleles were individually detected using allele specific antibodies that do not cross-react with the endogenous HLA-I alleles (Figure 2A). Thus, PAKC can be employed to investigate the properties of individual HLA-I alleles in the context of other (dys)functional APM component, which will improve our understanding of the association of the antigen presentation pathway with HLA-I associated diseases.

**Figure 2.**
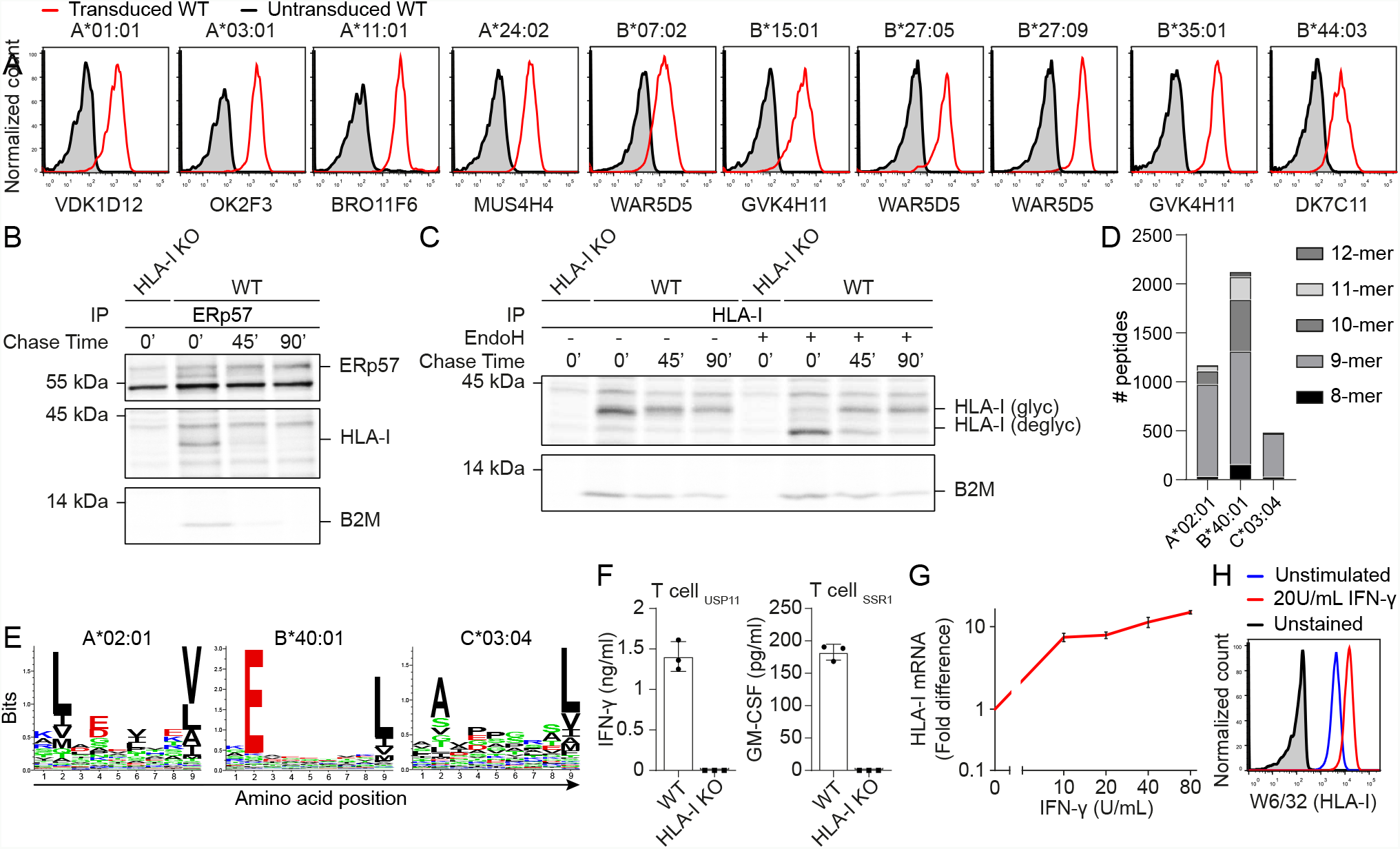
Research directions of PAKC. **(A)** HAP1 wild type cells transduced with different HLA-A and -B alleles and stained using indicated allele-specific antibodies (red), untransduced cells (grey). Gated on marker gene positive cells. **(B)**, **(C)** Analysis of the HLA-I maturation by pulse-chase in IFN-γ stimulated HAP1 wild type and HLA-I knockout cells metabolically labeled with [^35^S]-Met/Cys for 30 min and chased for 0, 45 or 90 min. **(B)** Co-immunoprecipitation of HLA-I using anti-ERp57 antibodies was visualized using phospho-imaging. **(C)** Co-immunoprecipitation using W6/32 for HLA-I followed or not by EndoH treatment. **(D)** 3,774 unique peptides were eluted from HAP1 wild type cells as identified by mass spectrometry and allocated to the allele with the highest predicted affinity using NetMHC4.0 with an affinity cutoff of 5 μM. **(E)** All identified 9-mer peptides were used to perform unsupervised sequence clustering and visualized by Seq2Logo to reveal common sequences for each HLA-I allele. **(F)** IFN-γ or GM-CSF secretion by HLA-A*02:01-restricted T cells specific for USP11- or SSR1-derived peptides after overnight co-culture with HAP1 wild type or HLA-I knockout cells. **(G)** Upregulation of HLA-I mRNA and **(H)** cell surface HLA-I after IFN-γ stimulation as assessed by qPCR and FACS, respectively. WT, wild type; KO, knockout.

After initial synthesis of the HLA-I heavy chain, the molecule matures through a process of folding, peptide loading and glycosylation events. We investigated whether PAKC allows for evaluation of individual APM component contributions to HLA-I maturation using pulse chase experiments. In wild type cells, ERp57 interacted only transiently with pulsed HLA-I, indicating that the HLA-I peptide loading largely occurred within 45 minutes (Figure 2B). The majority of HLA-I further matured within 90 minutes after pulse as detected by EndoH resistance of the *N*-glycan of immunoprecipitated HLA-I (Figure 2C). Such molecular studies will further catalyze new mechanistic and disease-related insights after overexpression of individual HLA-I alleles or mutant APM members in their respective knockout cells.

After two decades of algorithm development to predict the immunogenic HLA-I presented peptide repertoire, the scientific community is still facing major issues in the accuracy of these predictions. The contribution of several APM members to peptide selection has not been interrogated, mainly due to the lack of proper model cell lines. Although HAP1 cells have a lower HLA-I surface expression than the commonly used JY cell line, we investigated whether PAKC could be employed for HLA-I ligandome analyses on HAP1 wild type cells. Mass spectrometric analyses of eluted peptides identified 3774 peptides that were assigned to bind HLA-A*02:01, HLA-B*40:01 or HLA-C*03:04 based on peptide affinity prediction (Figure 2D). Sequence logos corresponding to the 9-mer peptides for HLA-A*02:01, HLA-B*40:01 and HLA-C*03:04 were in concordance with literature (Figure 2E)(16, 24, 25). Thus, PAKC is well-suited for high-resolution peptide elution studies which further benefit from the fact that PAKC is made on a single genetic and proteomic background. This property can be further exploited by studies to individual peptide processing and presentation which, besides targeted mass spectrometry, may be read out by peptide-specific T cell activation assays. This was confirmed by the cytokine production of two independent T cell clones recognizing endogenously derived peptides presented by HLA-A*02:01 on HAP1 wild type but not on control HLA-I knockout cells Figure 2F). Proteases other than ERAP1 described to affect peptide generation, including the proteasome, potentially have a profound effect on processing and presentation of individual peptides. These have not been included in the current PAKC, but it would be very useful to additionally include individual protease knockouts in the future. Thus, PAKC will contribute to a better understanding of the processing and HLA-I loading of peptides. The potential improvements of HLA-I ligandome prediction algorithms will advance translational research to infection, cancer and autoimmunity.

During infection, cancer and autoimmunity, antigen presentation is enhanced by locally produced cytokines including IFN-γ. The IFN-γ signaling pathway is intact in HAP1 cells(26). Our experiments now showed that induction of IFN-γ signaling upregulates HLA-I mRNA and surface protein levels (Figure 2E and F). These findings demonstrate that PAKC is highly suitable to study antigen presentation in an inflammatory context, for example in any of the above-mentioned research directions. Additionally, PAKC also supports research to non-classical HLA-I molecules, which are lowly expressed on HAP1 cells, but are expressed upon IFN-γ stimulation and exogenous peptide feeding. Finally, the haploid nature of HAP1 cells, and most of our clones, allows for powerful genome-wide haploid genetic screening to unravel novel antigen presentation biology(27).

In conclusion, we generated and validated a novel panel of ten cell lines, each knockout for an individual APM component which will accelerate antigen presentation research. We showed that this panel can be adopted into many different research lines as it is suitable for a wide variety of assays. As pointed out, PAKC will expand our knowledge by answering open questions in the field for which no good model systems were previously available(28). It will facilitate generation of new hypotheses, as well as revisiting and challenging published data that were generated in imperfect or inconsistent model systems over the years. PAKC is now available for the scientific community to enable unraveling of novel antigen presentation biology, some of which eventually may lead to new therapeutic approaches to treat infections, cancer and autoimmunity.

## Acknowledgments

We thank Dr. Ann-Charlott Schneider for performing qPCR and the Sanquin Research Facility for their assistance with FACS. Furthermore, we thank Dr. Sebastian Springer and Dr. Zeynep Hein (Jacobs University) for providing the HLA-B*27:05 and HLA-B*27:09 overexpression constructs, Dr. Marieke Griffioen (LUMC) for providing the SSR1 specific T cell clone, all other HLA-I overexpression constructs, as well as critically reading the manuscript, Dr. Mirjam Heemskerk (LUMC) for providing the USP11 specific T cell clone, Dr. Riccardo Mezzadra (The Netherlands Cancer Institute) for the B2M CRISPR/Cas9 construct and Dr. Jacques Neefjes (LUMC) for providing the HC10 antibodies.

---

**Supplemental Figure 1.**
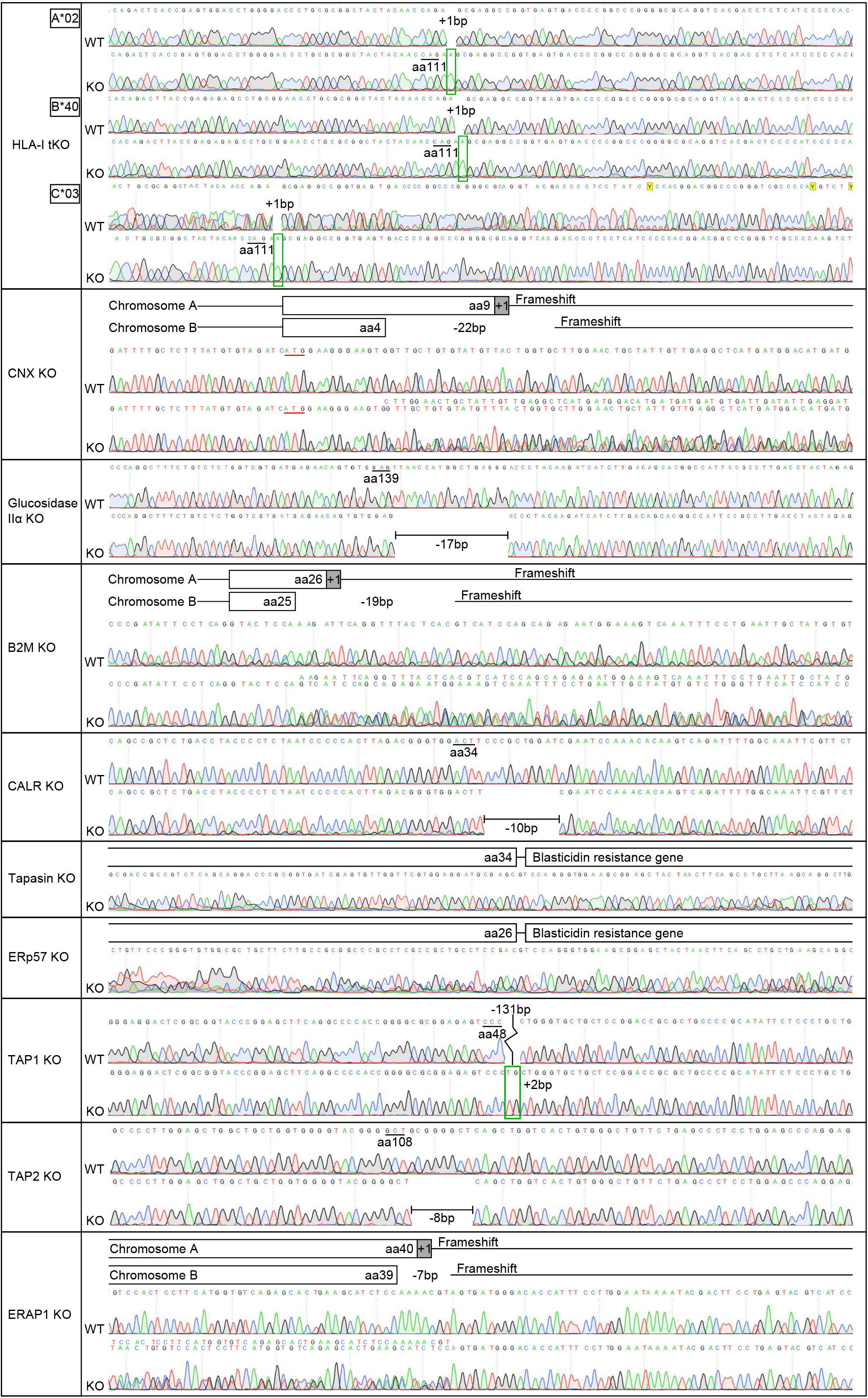
Sequencing details of PAKC. Sanger sequences around the Cas9 targeted site for each clonal cell line in PAKC. For each clone, wild type and knockout sequences are shown. For cases of blasticidin S-resistance gene insertion or mixed sequences due to diploidy of the cells, a graphical summary is shown above the sequences (upper sequence annotation refers to chromosome A and lower to chromosome B). The number of amino acids (aa) translated before the disrupting mutation are shown, underlined are translation start sites, green squares highlight base pair insertions. WT, wild type; KO, knockout.

**Supplemental Table I.**
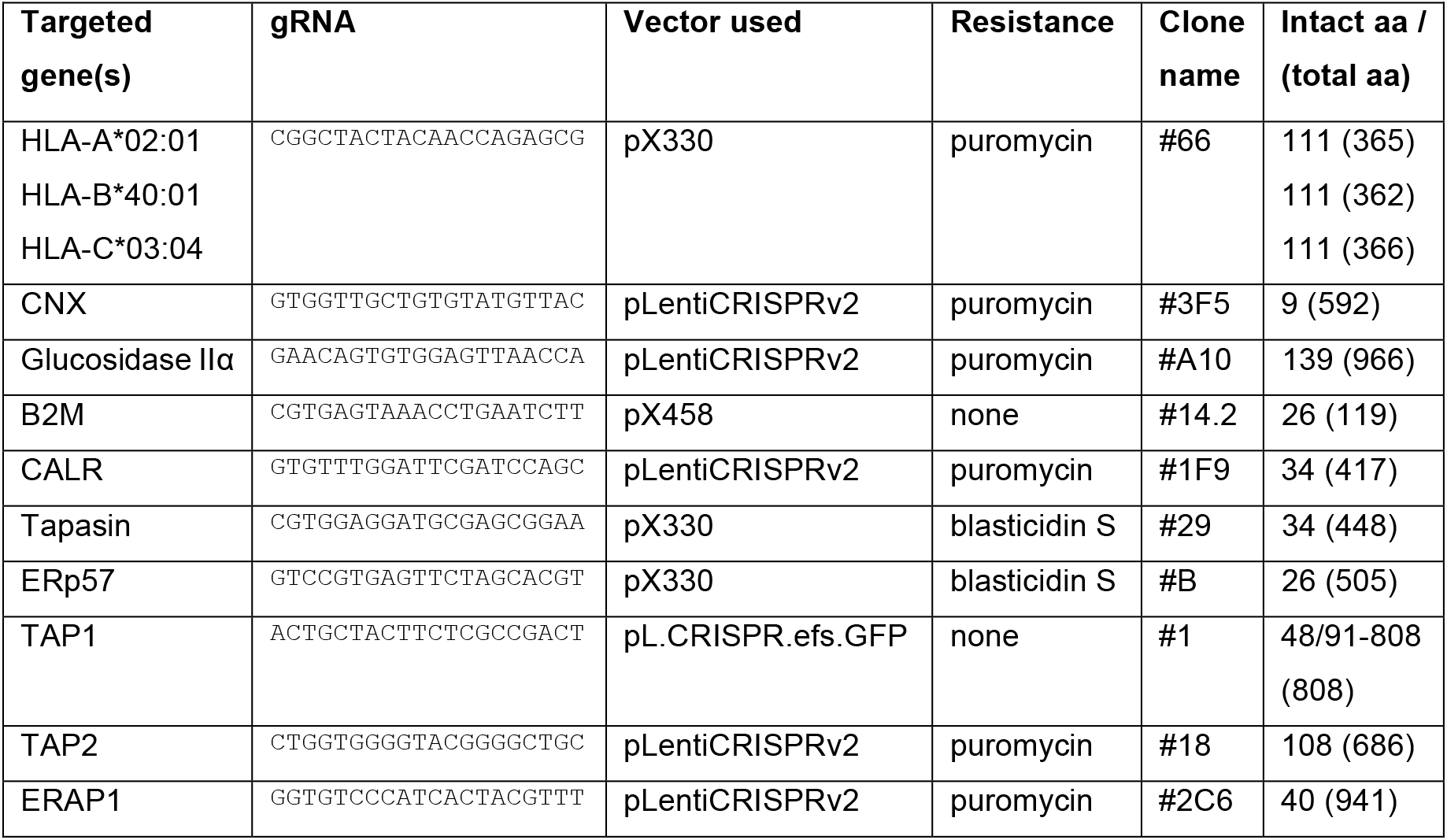
Summary of clonal cell lines of PAKC.

**Supplemental Table II.**
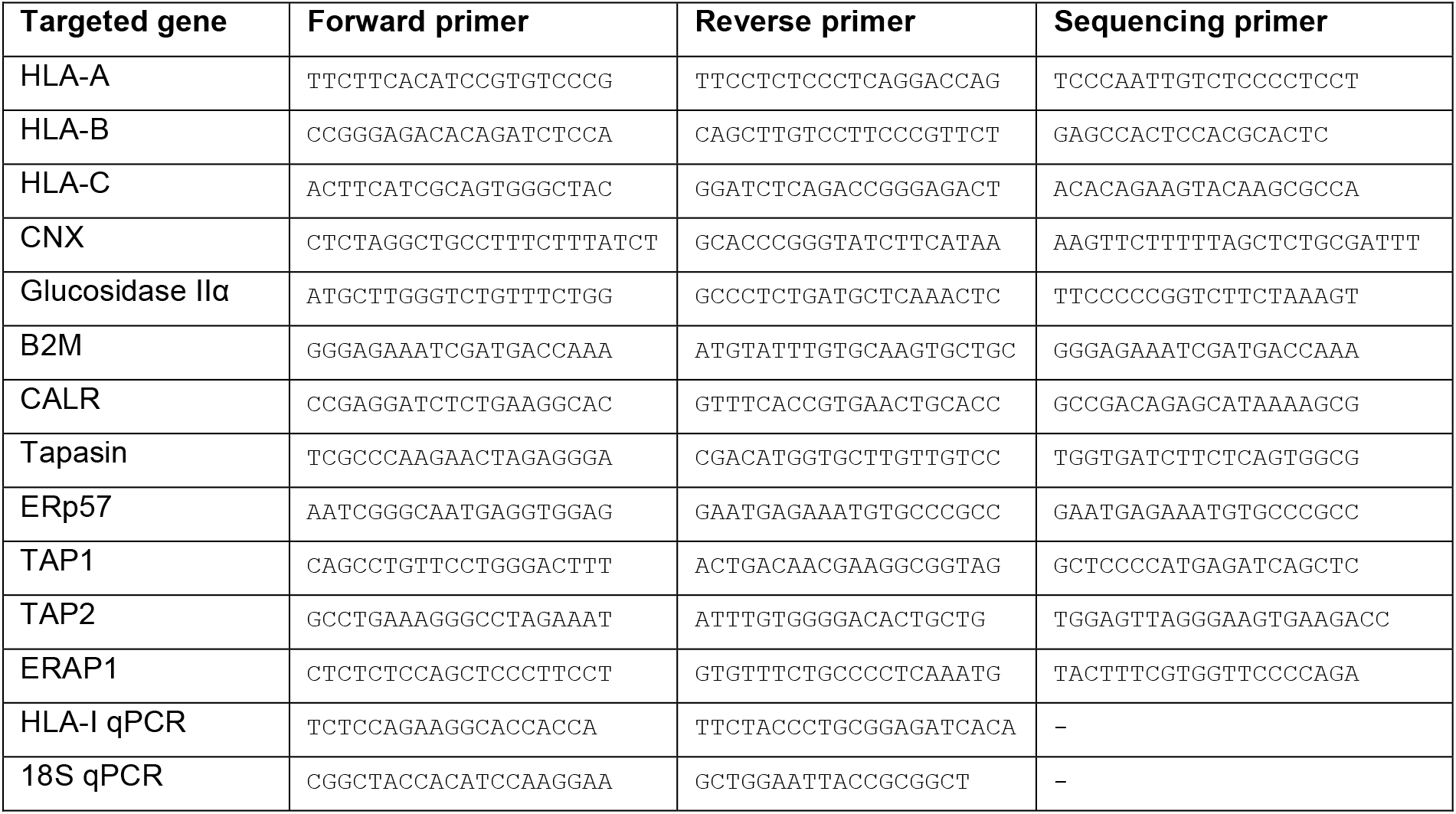
Primers used for PCR, qPCR and Sequencing.

**Supplemental Table III.** Antibody clones used for immunoblotting and flow cytometry.

| Antibody | Species | Used to Target | Manufacturer |
| --- | --- | --- | --- |
| Calnexin polyclonal | Rabbit | CNX | Stressgen<br>Biotechnologies<br>(SPA-865) |
| Calreticulin polyclonal | Rabbit | CALR | Sigma (C4606) |
| HC10 | Mouse | HLA-I | Kindly provided by<br>Dr. J. Neefjes |
| MaP.Erp57 | Mouse | ERp57 | Abcam |
| PaSta1 | Rat | tapasin | Self |
| 148.3 | Mouse | TAP1 | Self |
| AC-74 | Mouse | β-actin | Sigma-Aldrich |
| Anti-Mouse-HRP polyclonal | Goat | Mouse antibody | Sigma-Aldrich |
| Anti-Rabbit-HRP polyclonal | Goat | Rabbit antibody | Sigma-Aldrich |
| Anti-Rat-HRP polyclonal | Goat | Rat antibody | Sigma-Aldrich |
| W6/32-PerCP-eFluor710 | Mouse, IgG | HLA-A, -B, -C | eBioscience |
| VDK1D12 | Human, IgM | HLA-A*01:01 | Self |
| OK2F3 | Human, IgM | HLA-A*03:01 | Self |
| BRO11F6 | Human, IgG | HLA-A*11:01 | Self |
| MUS4H4 | Human, IgG | HLA-A*24:02 | Self |
| WAR5D5 | Human, IgG | HLA-B*07:02<br>HLA-B*27:05<br>HLA-B*27:09 | Self |
| GVK4H11 | Human, IgM | HLA-B*15:01<br>HLA-B*35:01 | Self |
| DK7C11 | Human, IgG | HLA-B*44:03 | Self |
| MH16-1 in-house conjugated to DL650 | Mouse, IgG | Human IgG | Sanquin |
| MHM-88-APC | Mouse, IgG | Human IgM | BioLegend |
| ME20.4-PECy7 | Mouse, IgG | Human NGFR | BioLegend |

## Notes

1 This work was supported by the Netherlands organization for scientific research (NWO-VENI 016.131.047; R.M.S.), KWF Alpe d’HuZes (Bas Mulder Award 2015-7982; R.M.S.), the Landsteiner Foundation for Blood Transfusion Research (LSBR fellowship 1842F; R.M.S.), the German Research Foundation (SFB 807 – Membrane Transport and Communication; R.T.) and an ERC Advanced Grant (EditMHC; R.T.).

